# Hyaluronic acid-b-polylactic acid polymersomes facilitate CD44-mediated delivery of doxorubicin to glioblastoma in vitro

**DOI:** 10.64898/2026.05.26.727934

**Authors:** Apoorvi Chaudhri, Molli Garifo, Pranavi Thatravarthi, Torrick Fletcher, Jessica Larsen

## Abstract

Glioblastoma represents a highly aggressive brain tumor with low survival and no response to chemotherapy and radiation therapy. Temozolomide, the current standard of care chemotherapy, improves patient survival by only about 6 months because of several resistance mechanisms, including unmethylated MGMT, which enables repair of chemotherapy-induced DNA damage. Thus, additional treatments strategies are necessary to investigate efficient responses towards glioblastoma. Doxorubicin (DOX) is a chemotherapeutic agent that is independent of MGMT methylation and instead works through inhibition of topoisomerase (TOPO) II, an enzyme necessary for DNA replication of the tumor. The inability of doxorubicin to cross the blood-brain barrier (BBB) precludes its use in glioblastoma. Polymersome nanoparticles have the potential to transport agents across the BBB. Here, we develop hyaluronic acid-b-polylactic acid (HA-PLA) polymeric nanoparticles called polymersomes, encapsulate them with DOX and investigate the ability of our system to induce apoptosis in a human glioblastoma cell line. The HA-PLA polymersomes show specificity and receptor-mediated endocytosis towards CD44-positive U87 glioblastoma cells due to the natural affinity of HA (hyaluronic acid) to CD44. Our HLA-PLA-DOX system promotes apoptosis of glioblastoma through inhibition of topoisomerase (TOPO) II. Thus, our system could allow tumor specificity through HA-CD44 affinity and slow drug release through pH sensitivity of PLA in the acidic tumor microenvironment.

## Introduction

Glioblastoma is an aggressive tumor with a low survival rate of only 9 months^1^. Temozolomide, the standard chemotherapy for glioblastoma, increases the survival rate to only 15 months^2^. Resistance towards temozolomide includes unmethylation or overactivation of O6-methylguanine-methyltransferase (MGMT) that prevents DNA damage and cellular apoptosis^3^. Doxorubicin is a chemotherapeutic agent that instead promotes tumor cell toxicity through the inhibition of topoisomerase II enzyme (TOPOII)^4^. Tumor cells are dependent on TOPOII for DNA replication and cell proliferation, a key event in cancer growth^5^. Doxorubicin is a standard chemotherapy in multiple cancer types, including breast cancer, bladder cancer and non-small lung cancer^6^. A major challenge to the use of doxorobucin in glioblastoma treatment is its penetration across the blood-brain barrier (BBB). It is hydrophilic nature, has a high molecular weight and is a substrate for transporter proteins that result in low concentration in the brain^7^. This inability of doxorubicin to reach the glioblastoma tumor has prevented it from being an effective treatment strategy in this disease. Nanoparticles can act as delivery vehicles to effectively transport drugs across the BBB in multiple settings^8,9^. Polymersomes are nano-vesicles that are formed via self-assembly of amphiphilic block copolymers^10,11^. HA-PLA (hyaluronic acid-b-polylactic acid) derived polymersomes are formed from a block copolymer of hyaluronic acid and polylactic acid that self-assemble into vesicles based on their molecular weights^12,13^. HA has a natural affinity to CD44 that facilitates nanoparticle internalization through receptor-mediated endocytosis in CD44+ tumor cells^14^. Multiple tumor models have used HA nanoparticles to deliver agents in CD44-expressing tumor cells^15,16^. Further, PLA nanoparticles are engineered to release drugs in acidic tumor microenvironments via acid-mediated hydrolysis^17^. Specifically, Deng et al. observed lower lung tumor metastasis in a 4T1 breast tumor xenograft model when treated with RGD-peptide conjugated HA-b-PLA-DOX. In mice, DOX had higher blood circulation time when loaded in these nanoparticles^18^. While HA-PLA polymersomes with doxorubicin have been tested in breast tumor models, less is known about their efficacy in glioblastoma. We hypothesize that DOX will be an effective therapy in glioblastoma when delivered through a HA-PLA based polymersome. Our HA-PLA polymersome with its hydrophobic PLA membrane encapsulates the hydrophilic DOX. Dynamic light scattering shows the HA-PLA-DOX- loaded polymersomes vary in diameter based on DOX loaded content but maintain a negative surface charge. HA-PLA polymersome loaded with FITC-BSA shows CD44 receptor mediated endocytosis and internalization in CD44+ U87 cells but not CD44- Jurkat cells. Further, HA-PLA-DOX shows toxicity against CD44+ U87 glioblastoma cells, as evidenced by reduced tumor cell proliferation through DOX-mediated inhibition of TOPOII activity. Together, we propose our HA-PLA-DOX system as a superior therapeutic strategy for efficient delivery of doxorubicin to CD44+ glioblastoma tumor cells.

## Experimental

### Materials

HA, with an average molecular weight of 7 kDa, was purchased from Creative PEGWorks (Durham, NC, USA). N-hydroxysuccinimide (NHS) functionalized PLA (PLA-NHS) was purchased from PolySciTech, Akinalytics (West Lafayette, IN, USA) at an average molecular weight of 15 kDa. 1,4-diaminobutane was purchased from Thermo Fisher Scientific (Waltham, MA, USA). Sodium cyanoborohydride (NaCNBH3, 99.2%) was purchased from Chem-Impex International (Wood Dale, IL, USA). N, N-di-isopropyl ethylamine (DIPEA) was purchased from Thermo Fisher Scientific (Waltham, MA, USA). DOX was purchased from APExBIO (MW ∼0.5 kDa, Houston, TX, USA). Acetate buffer, dimethyl sulfoxide (DMSO), mannitol, sodium chloride, phosphate-buffered saline (PBS), 4-(2-hydroxyethyl) piperazine-1-ethane-sulfonic acid (HEPES) buffer were obtained from Sigma-Aldrich (St. Louis, MO, USA).

U87-MG cells were purchased from ATCC (Manassas, VA, USA) to perform *in vitro* studies on malignant glioma cells. Standard cell media comprising of Minimum Essential Medium Eagle with Earle’s salts, non-essential amino acids, L-glutamine, sodium bicarbonate, fetal bovine serum, and 1x penicillin-streptomycin solution were purchased from Sigma-Aldrich (St. Louis, MO, USA). Passages were performed using 0.25% trypsin (Corning, Inc., NY, USA). CellTriter 96 Aqueous One Solution Cell Proliferation (MTS) Assay was purchased from Promega (Madison, WI, USA). TOPO II Assay Kit was purchased from TopoGEN (Buena Vista, CO, USA). Quick-DNA Miniprep was purchased from Zymo Research (Tustin, CA, USA). Gel electrophoresis was performed using TAE buffer, synthesized following standard recipe using Tris base (≥99.8%) from Bio-Rad Laboratories (Hercules, CA, USA), glacial acetic acid (≥99%) from VWR (Radnor, PA, USA), and EDTA (≥99%) from Alfa Aesar (Ward Hill, MA, USA) and agarose from Thermo Fisher Scientific (Waltham, MA, USA). Ethidium bromide (EB) for gel staining was purchased from Thermo Fisher Scientific (Waltham, MA, USA). Albumin from bovine serum with fluorescein conjugate (FITC-BSA, MW ∼66 kDa) was purchased from Sigma-Aldrich (St. Louis, MO, USA).

### HA-PLA synthesis

Hyaluronic acid-β-polylactic acid (HA-PLA) copolymer was synthesized in a two-step conjugation method as described in a previous study (15). The HA was selected to have a molecular weight of 7 kDa based on previous work performed in our laboratory. When enzyme-responsive polymersomes were made from HA 7 kDa and 15 kDa PLA, they exhibited a small nanoparticle diameter of 76.5 ± 7.7 nm, high loading capabilities using our osmotic pressure drive loading approach, and show the greatest release rate variance between physiologic conditions and pathologic conditions^12^.

### Amination reaction of hyaluronic acid

A terminal reduction amination reaction was performed to functionalize HA for the conjugation step. In detail, 0.1 g of HA was dissolved in excess 0.1 M acetate buffer (pH 5.4) before adding 1 mL 1,4-diaminobutane dropwise under magnetic stirring. The amination reaction was ongoing for 24 hours at 50° C. NaCNBH_3_ (0.2 g) was added as a standard reducing agent with stirring at 50°C. An additional 0.1 g of NaCNBH_3_ was added 24 hours after the first addition to ensure the reduction was complete. The mixture was dialyzed against deionized water for 72 hours to remove all excess unreacted 1,4-diaminobutane and NaCNBH_3_ using a 3.5 kDa molecular weight cut-off (MWCO) dialysis cassette (Spectrum Labs, CA, USA). The dialyzed mixture was then filtered using a 0.45 μm syringe filter (Millipore, MA, USA) before lyophilizing to obtain aminated HA in solid form.

### Conjugation of aminated hyaluronic acid and N-hydroxysuccinimide functionalized polylactic acid

The conjugation of PLA-NHS and aminated HA was performed using the NHS ester-amine coupling reaction^12^. These procedures involved 0.3 g of PLA-NHS dissolved in 10 mL DMSO, and 0.2 g (stoichiometric excess) of animated HA added with 15 μL of DIPEA, which served as a catalyst for the coupling reaction. The reaction flask was housed in a sand bath and left to react for 48 hours, prior to heating at 50°C inside a glove box to ensure the reaction occured under nitrogen. To remove all excess unreacted reactants, the reaction mixture was dialyzed against deionized water for 72 hours using a 10 kDa MWCO dialysis cassette (Spectrum Labs, CA, USA). The dialyzed mixture was then filtered using a 0.45 μm syringe filter before lyophilizing to obtain HA-PLA in solid form.

### HA-PLA polymersome synthesis

The copolymer was used for polymersome synthesis via solvent injection. Briefly, HA-PLA was dissolved in DMSO at a concentration of 2.4mg/100μL. An aqueous solution of 8 wt. % mannitol in deionized (DI) water was used to enhance the polymersome stability during lyophilization. The polymer solution was injected into the aqueous solution using a syringe pump at a rate of 20 μL/min through a 20-gauge needle following previously established protocols from our laboratory^19^. Synthesized polymersomes were filtered through a 0.45 μm syringe filter before characterization.

### HA-PLA Polymersome characterization

Dynamic light scattering (DLS, Zetasizer Nano ZS90, Malvern Instruments, Malvern, UK) determined the particle size and ζ-potential of the polymersomes. One mL of the filtered polymersomes was used for size measurements. Sodium chloride was added to 1 mL of filtered polymersomes to obtain a concentration of 100 μM for ζ-potential measurements, as ions/counterions in solution are required to enable this measurement. The remaining polymersome solution was lyophilized for future characterization following previously established protocols^20^. Polymersomes were prepared for transmission electron microscopy at a concentration of 1 mg/mL. Phosphotungstic acid at 1% was applied for negative staining. TEM images were taken on a Hitachi 7830 UHR transmission electron microscope (Tokyo, Japan) at 120 kV.

### Payload encapsulation in HA-PLA polymersome

Polymersomes were actively loaded with FITC-BSA or DOX at varying concentrations. Encapsulation efficiency (EE) and release profiles were calculated on a mass basis using UV-Vis analyses and appropriate internal calibration curves.Appropriate wavelength for UV-Vis analysis, a spectral scan was performed on DOX in DI water and pH 7.4 and 6.8 HEPES buffer at concentrations ranging from 0 to 24 μg/mL (Table S1). Lyophilized polymersomes were dissolved into deioinized water at a concentration of 10mg/mL containing 2 mg/mL FITC-BSA or known concentrations of DOX (3, 6, 12 or 24 μg/mL, C_0_). 500 μL of sample was transferred to 100 kDa microcentrifuge dialysis tubes with 750 μL deionized water as the dialysate. The tubes were placed in a tube rack on a shaker plate for constant movement at room temperature. Every hour for four-six hours, the dialysate was analyzed using UV-Vis spectrophotometer to quantify the mass of DOX released (C_r_). The dialysate was replaced after every sample was collected to maintain a concentration gradient. Any DOX not released through dialysis and not washed away from the surrounding fluid and the polymersome surface, was assumed to be encapsulated. EE was calculated using the following equation:

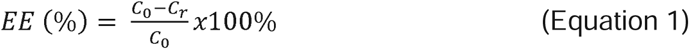

### Doxorubicin release from HA-PLA polymersomes

DOX-loaded polymersomes were concentrated using 100 kDa MWCO spin-down columns (Millipore, MA, USA). The concentrated, DOX-loaded polymersomes were resuspended in 100 μL pH 7.4 and pH 6.8 HEPES buffer to model both normal tissue and tumor microenvironments, respectively. The samples were transferred into 100 kDa mini dialysis tubes to allow transport of DOX but not the polymersomes, with 1.2 ml corresponding buffer as the dialysate. The tubes were placed in a tube rack on a shaker plate in an incubator at 37°C to simulate physiological conditions. The dialysate was sampled over a minimum of 60 hours and analyzed using UV-Vis spectroscopy to quantify the mass of DOX released. Calibration curves are used to quantify the mass and percentage of DOX released.

### Cells and cell culture

U87-MG cells were maintained in EMEM (Minimum Essential Medium Eagle with Earle’s salts) with 10% heat inactivated FCS, 1% glutamax, and 1% pen-strep in 5% CO_2_. U87 cells were obtained from ATCC. Jurkat-NFAT cells were obtained from BPS technologies and maintained in BPS5 media; RPMI 1640 (Roswell Park Memorial Institute) with 10% heat inactivated FCS, 1% glutamax, 1% pen-strep, 1% HEPES, 1% pyruvate and G418 (500 µg/ml) in 5% CO2. For polymersome uptake assays, jurkat NFAT cells were plated in BPS4 media; RPMI 1640 (Roswell Park Memorial Institute) with 10% heat inactivated FCS, 1% glutamax, 1% pen-strep, 1% HEPES, 1% pyruvate in 5% CO2. These cell lines tested negative for mycoplasma. The U87 and Jurkat NFAT cell lines were assayed with anti-human CD44 antibody (Biolegend clone 1M7) using flow cytometry. The corresponding isotype was used as control.

### HA-PLA polymersome cellular uptake

Polymersomes were loaded with FITC-BSA (MW ∼66 kDa, Sigma Aldrich) to study uptake behavior, as DOX would kill the cells. U87-MG cells or Jurkat-NFAT were seeded in a 96-well plate at 75,000 cells per well. After 24 hours, media was replaced and the wells were treated with the following groups: media (control), free FITC-BSA, or HA-PLA polymersome (2.5 mg), loaded at FITC-BSA concentration of 2 μg/mg polymersome. For Jurkat cells, the wells were pre-treated with poly-lysine to promote adherence. Treated cells were incubated for 4 hours at 37 LC or 4 LC. Upon incubation, cells were harvested and acquired on Cytek Aurora (CA, USA) flow cytometer to quantify the uptake of HA-PLA polymersomes. Data was analyzed using flow-jo software version 10. Confocal microscopy was also used to visualize cellular uptake. Briefly, U87 or Jurkat cells cells were treated with conditions as described above for 4 hours at either temperature. After incubation, cells were fixed with ice-cold 4% PFA for 10 min on ice. On fixation, cells were washed with PBS, chambers were removed, and Vectashield antifade mounting medium with DAPI (Vector Laboratories H-1200-10) was added to the slides. The slides were sealed and incubated at least overnight at 4 LC before microscopy. Images were acquired using Leica Stellaris 5 confocal microscope. For competitive uptake experiments, varying amounts of HA-PLA-FITC-BSA and 10 mg/mL free HA were added during a 4-hour incubation at 37 LC. Uptake was analyzed by flow cytometry, with gates set to those of untreated controls.

### Calculation of LC-50 for free DOX

A dose escalation study was performed to determine which DOX concentrations should be encapsulated to achieve effective chemotherapeutic activity. U87-MG cells were seeded in a 96-well plate and allowed to sit overnight. Following, cells were treated with free DOX added to fresh growth media at final DOX concentrations 0-24 μg/mL. At days 1-5 post incubation, cell death was evaluated using the MTS proliferation assay. The MTS reagent was added to each well at 10% working volume and incubated for 4 hours. The absorbance was measured at 490 nm using spectrophotometer (UV-Vis, BioTek Synergy H1, Aligent Technologies; CA, USA). LC50 values were calculated by fitting to a curve in GraphPad Prism.

### MTS cell viability assay

Viability studies were performed on empty and loaded polymersomes. U87-MG cells were seeded in a 96-well plate at 10,000 cells per well. When cells reached 70% confluency, cells were treated with 200 μL fresh media, 6 μg/mL and 24 μg/mL free DOX, and 6 and 24 μg/mL DOX-loaded polymersome. The viability studies were performed using an MTS proliferation assay 4 and 5 days after dosing as described in Ma et al^21^.

### Topoisomerase II assay

Cells were dosed with U87-MG cells as in viability studies and collected after five days of incubation. DNA was extracted using the Zymo Quick-DNA Miniprep Kit. The TOPO II assay kit (TopoGEN Kit, TG1001) was used to detect the topoisomerase enzyme activity. The assay was performed based on the manufacturer’s instructions. Briefly, the enzymatic reaction mixture contained 4 μL of 5X Complete Reaction Buffer, 200 ng kDNA substrate, 2 IU of TOPO II and 80-120 ng DNA extract from the treated cells. The mixture was incubated at 37C for 30 mins followed by addition of 5x Stop Buffer. The reaction products were run in a 1% agarose gel.

### Statistics

Results were graphed as mean with SEM (standard error of mean). The statistical analysis was performed using GraphPad Prism version 10 using 2 Way ANOVA with multiple comparisons and Tukey correction for multiple comparisons.

## Results

### HA-PLA polymersomes can be synthesized with uniform diameters

We synthesized HA-PLA polymersomes using a solvent injection method as previously described (18). The polymersomes were characterized by dynamic light scattering and transmission electron microscopy (Figure 1A, C, D). Empty HA-PLA polymersomes without DOX were included as a control. Empty HA-PLA polymersomes form at diameters around 100 nm with monodisperse polydispersity indices (PDI), while the addition of DOX appears to increase the polymersome diameter and the PDI. This increase in PDI is likely due to the innate fluorescence and colorimetric nature of DOX^22^. DLS utilizes the material’s refractive index to determine size and PDI, and DOX naturally fluoresces red, rendering the solution colorimetric which can be disruptive to sample measurement^23^. Notably, TEM images identified relatively monodisperse particle diameters with DOX loading (Figure 1D). Overall, polymersomes regardless of DOX loading presented with an average size between 100-150 nm and a negative charge, suggesting effective uptake in the cells. A slightly negative surface charge can facilitate cellular uptake^24^, specifically in malignant cell lines^25^.

**Figure 1.**
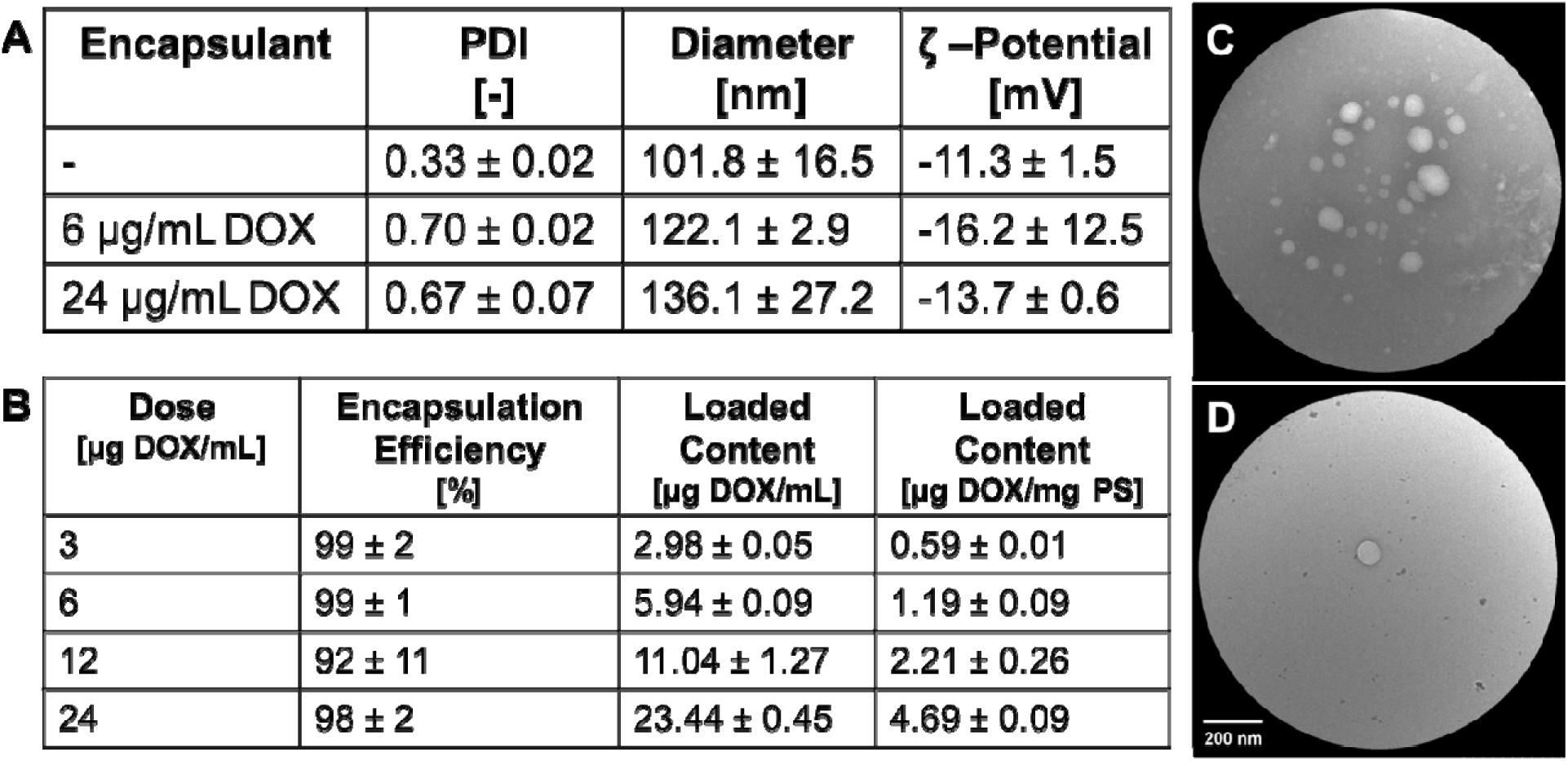
A. Polymersome characterization via dynamic light scattering with and without DOX encapsulation, B. Characterization of the encapsulation of DOX at various concentrations. C. Representative TEM image of empty HAPLA polymersomes. D. Representative TEM image of DOX-loaded HA-PLA polymersomes (6 µg/mL).

### Doxorubicin can be encapsulated at high efficiencies and with high loading content into HA-PLA polymersomes

HA-PLA polymersomes were actively loaded with varying concentrations of DOX to ensure a therapeutically significant amount could be encapsulated and to ensure a maximum encapsulation was reached. 3, 6, 12, and 24 μg DOX were added into a 1 mL sample containing 5 mg of lyophilized polymersomes to determine encapsulation efficiency. All formulations achieved over 98% EE on average, aside from 12 μg, which had very large deviation within replicates (Figure 1B); however, all EE values were statistically similar to one another. The high EE is attributed to the addition of mannitol in the solvent during synthesis, as it acts as an osmotic agent^26^. The solvent injection method produced a range in polymersome sizes (Figure 1A), highlighted by the increasing standard deviation in diameters. This deviation in size could explain the slight deviation in EE. Loaded content was also calculated on a per mL and per mg polymersome basis, which provided useful information for designing in vitro and in vivo studies, as it allows for equal DOX dosing across various formulations. As expected, based on consistent EE, the loaded content increased proportionally with the amount of DOX initially loaded.

### Doxorubicin release is driven by diffusion and polymersome degradation

Loaded HA-PLA polymersomes were incubated in physiologic and tumorigenic mimetic buffers. The acidic condition was 0.1M HEPES Buffer at pH 6.85 to best represent the GBM tumor microenvironment^27,28^, whereas the neutral condition was 0.1M HEPES Buffer at pH 7.4 to represent the healthy brain environment. The study was performed for at least 60 hours. Figure 2 illustrates the release profile for both acidic and neutral conditions. The lower concentrations of 3 (Figure 2A), 6 (Figure 2B), and 12 (Figure 2C) μg DOX/mL resulted in little to no statistical significance in DOX released from HA-PLA polymersomes regardless of pH, likely due to the low overall DOX-loaded content; however, at 24 μg DOX/mL, statistically different release profiles were present between the buffer conditions after 36 hours (Figure 2D). The low overall loaded content of DOX when only 3 µg/mL is added and the dramatic deviations observed in the loaded content of DOX when 12 µg/mL is added (Figure 1B) are likely contributing to the large error bars observed in these release studies. For these reasons, further analysis included only HA-PLA polymersomes encapsulated with either 6 or 24 µg/mL.

**Figure 2.**
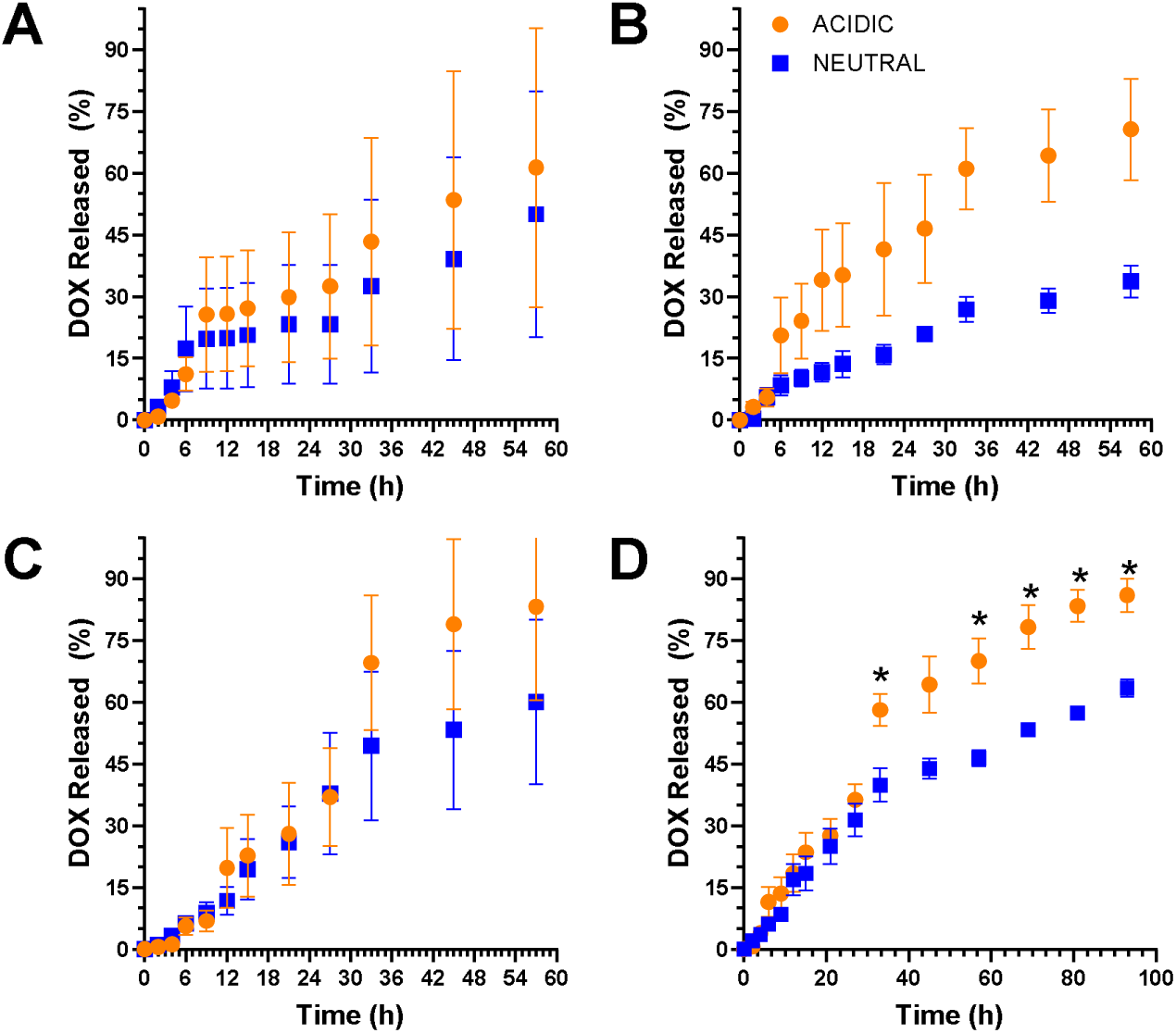
Release of DOX from HA-PLA polymersomes after various starting concentrations of DOX. (A) 3 µg/mL, (B) 6 µg/mL, (C) 12 µg/mL, and (D) 24 µg/mL

The lack of burst release at early time points suggests that DOX is encapsulated within the polymersome, as described by Wang et al^29^. In the first few hours of release, regardless of the amount of DOX loaded, the release profile was similar across pH values, suggesting that early DOX release from polymersomes was diffusion-driven and required DOX to transport through the PLA membrane. This prolonged release of DOX is expected; Oz et al. observed a similar profile from PLA-based polymersomes in pH 7.4 PBS buffer, with a maximum cumulative DOX release of 70%^30^. Multiple studies on the DOX release profile from PLA-based nanoparticles indicate an increase in release rate in more acidic environments, resulting from PLA degradation via acidic hydrolysis of ester bonds^31–33^, which was also observed in our system.

The data was then fit to the Korsmeyer-Peppas model (Equation 2) to establish the exponential relationship between release and time^34^.

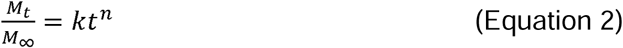

M_t_ represents the cumulative drug released at time t, M∞ represents the cumulative amount of drug released at infinite time, k is the release rate, and n represents the release exponent relating to the drug release mechanism. When n = 0.5, the release kinetics are described as Fickian drug diffusion^35,36^. When 0.5<n<1, the model assumes non-Fickian release, and release can be driven by other properties like polymer relaxation/degradation. For our system, we expect to see 0.5<n<1 as initial drug release should resemble Fickian diffusion through the HA-PLA membrane, with a shift to polymersome degradation-drive release over time. As a result, n should be representative of non-Fickian diffusion overall, and k should be larger in acidic conditions as compared to neutral conditions to represent an increased release rate. The 6 µg/mL DOX formulation shows large differences in the release rate between acidic and neutral environments (Figure 3A), suggesting PLA hydrolysis-driven DOX release that is more rapid in the acidic environment. Although the overall release rates when using the 24 µg/mL DOX HA-PLA formulation appear to be similar across the entire timeline of the study, post-24-hour release rates begin to differ, with higher release rates in an acidic environment. This suggests that the first 24 hours of DOX release are driven by diffusion, while after 24 hours PLA hydrolysis-driven release becomes the driver (Figure 3B).

**Figure 3.**
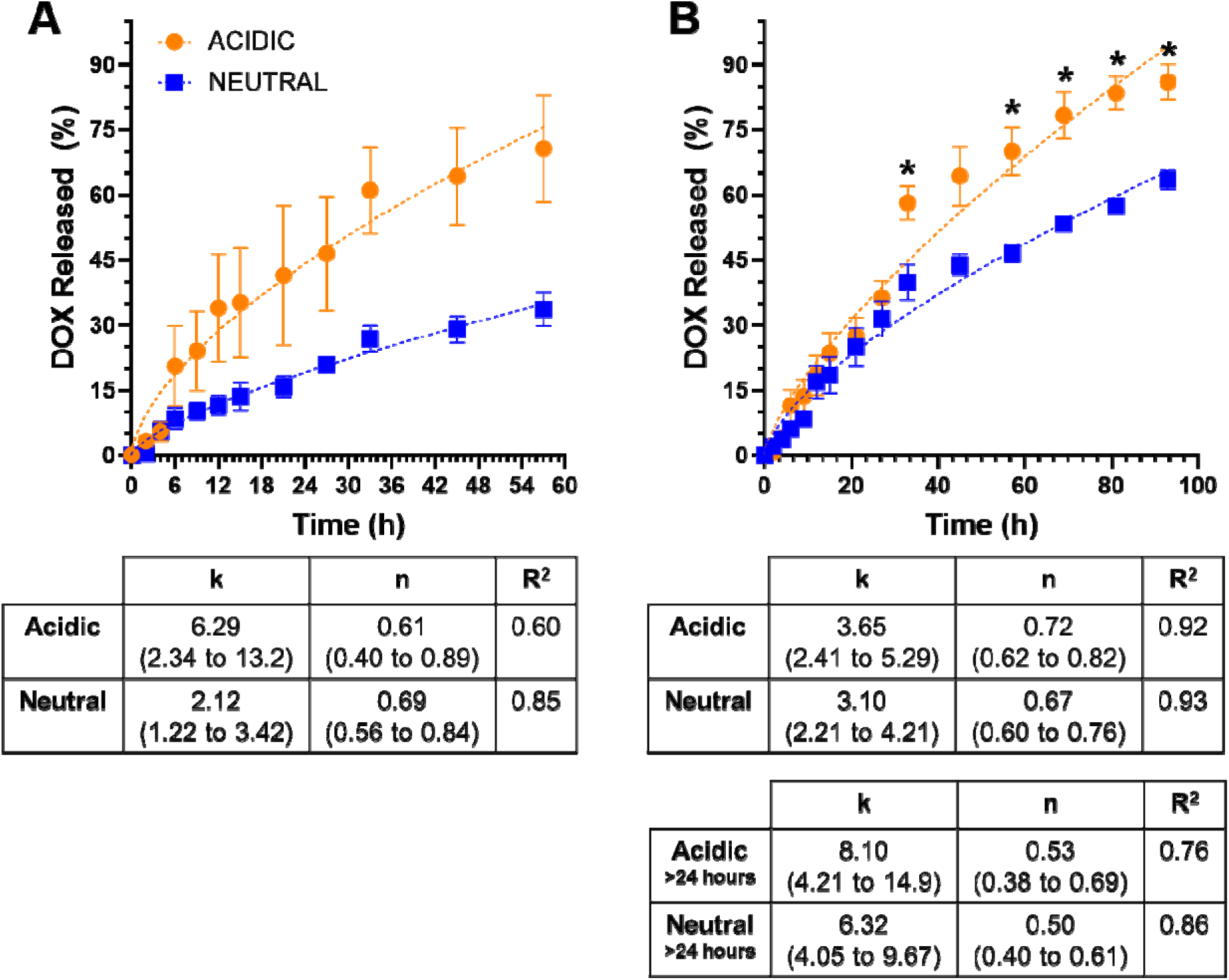
Release of DOX from HA-PLA polymersomes loaded with (A) 6 µg/mL or (B) 24 µg/mL is fit to the Korsemeyer-Peppas equation to highlight differences in release rates and release orders observed in each formulation.

### HA-PLA polymersomes are internalized by CD44 receptor-mediated endocytosis on U87 glioblastoma cells

After confirming that U87 cells present with the CD44 receptor, while Jurkat cells do not (Figure S1), we loaded HA-PLA polymersomes with FITC-BSA to determine cellular uptake capability of HA-PLA polymersomes. It is important to note that uptake experiments using DOX-loaded polymersomes would not have been effective, as DOX can kill the cells. Similarly, 10 µL of 2 mg/mL of FITC-BSA was added to 10mg lyophilized polymersomes and the encapsulation efficiency was determined to be 83.9 ± 6.7% with a loaded content of 1.68 ± 0.13 µg/mg HA-PLA, which is similar to our previous findings^12^. To determine cellular uptake, U87 cells were treated with 2.5 mg HA-PLA polymersomes loaded with FITC-BSA and incubated at 37 LC for 4 hours. Polymersome uptake was examined by flow cytometry. HA-PLA-FITC-BSA showed approximately 41% cellular uptake, indicating successful internalization (Figure 4A, C) with a high mean fluorescence intensity (MFI) (Figure 4D). Confocal microscopy also showed internalization with HA-PLA polymersomes loaded with FITC-BSA and free-FITC-BSA, while no treatment did not show any green fluorescence, as expected (Figure 4E). HA-PLA-FITC-BSA uptake in CD44-expressing U87 cells was compared with that in Jurkat cells lacking CD44 expression^37^ under the same temperature conditions and doses. Notably, uptake was minimal, with only around 2% of cells positive for polymersomes (Figure 4B,C) and extremely low MFI (Figure 4D).

**Figure 4.**
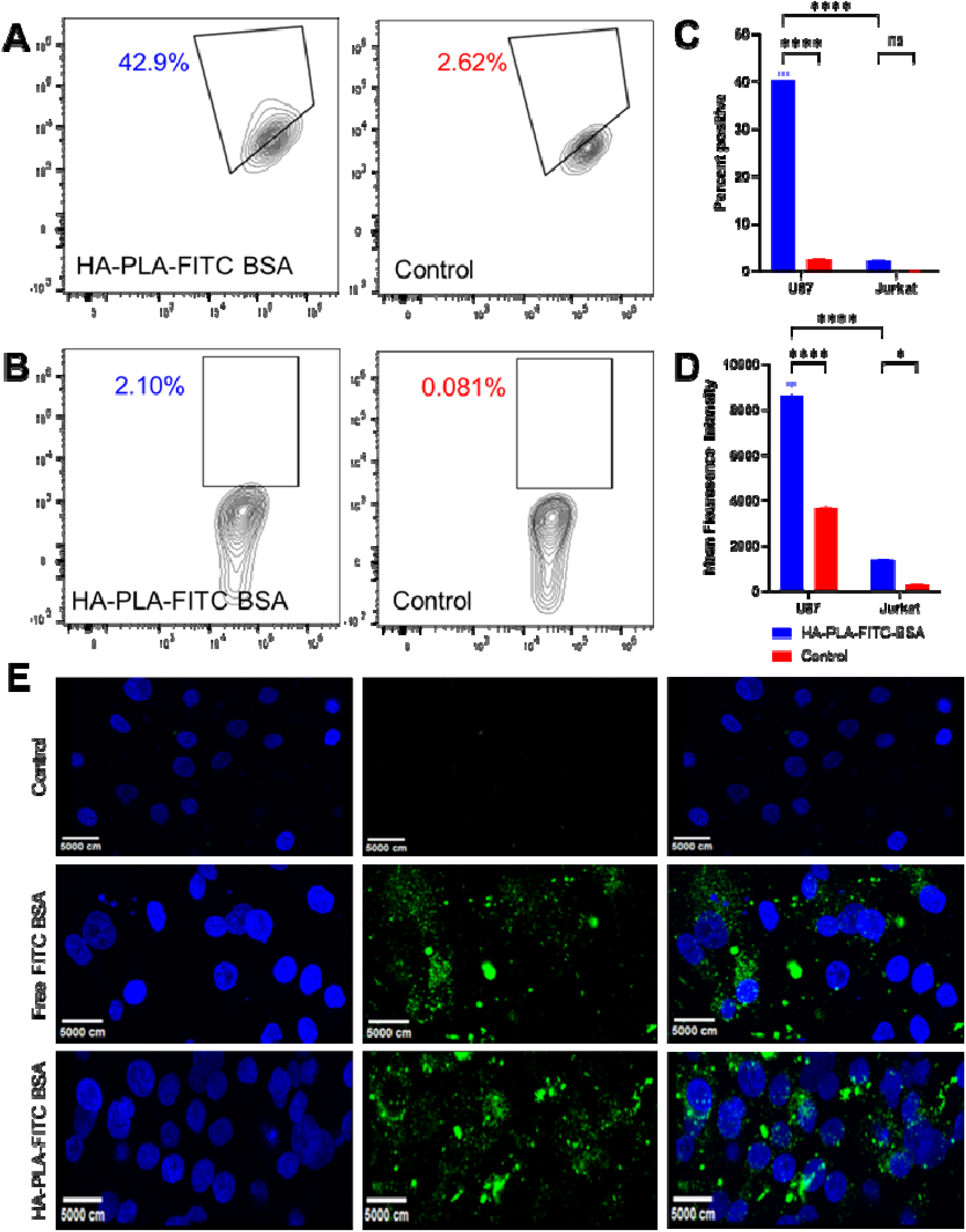
Example flow cytometry plots from HA-PLA-FITC BSA incubation with (A) U87 cells and (B) Jurkat cells at 37 □C. Flow cytometry data was analyzed over triplicates to calculate (A) the percent of cells positive for FITC-BSA and (B) the mean fluorescence intensity. (E) Confocal imaging confirms HA-PLA-FITC-BSA is taken up by U87 cells.* p<0.05, ** p<0.01, *** p<0.001, **** p<0.0001

Interestingly, treatment at 4 LC greatly reduced the uptake capacity of HA-PLA-FITC-BSA polymersomes in U87 cells from approximately 41% to approximately 12% (Figure 5A, C) and around half the MFI (Figure 5D). Changes in uptake between 37 LC and 4 LC were not as apparent with Jurkat cells with similar positive cells and MFI (Figure 5B, C,D). Simultaneously, competitive uptake experiments changed the HA-PLA-FITC-BSA uptake behavior. While the average uptake of the polymersome is between 60-80% when in the competitive environment, the MFI was dramatically reduced, which indicated fewer polymersomes entered each cell (Figure 6). Previous studies performed by Deng et al. utilized the addition of free HA within the cell culture media to evaluate CD44-HA relationship for cellular uptake. The addition of free HA provided a competitive environment for the HA-PLA polymersomes to bind to CD44 receptors on the surface of 4T1 cells, a metastatic breast cancer cellular model^18^. The drastic reduction in MFI was similar to what was observed in this study, where a 14% reduction in MFI was observed after a one-hour incubation. We hypothesize the difference between our findings is a result of the significant increase in CD44 expression in GBM (98%) as compared to breast cancer cells (∼30%) and a longer incubation period^38^. Together, our findings strongly suggest that the mechanism behind polymersome internalization was CD44 receptor-mediated endocytosis.

**Figure 5.**
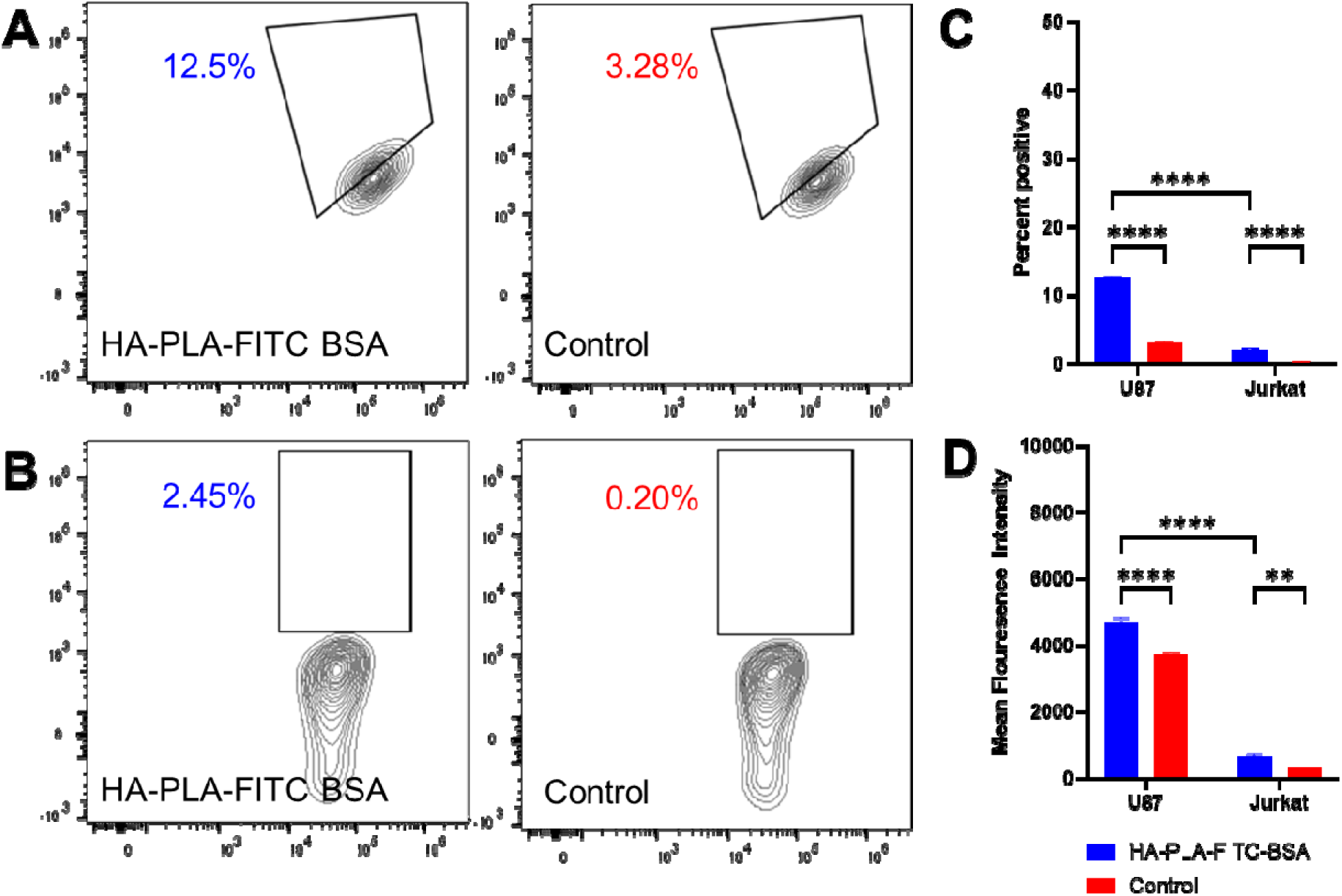
Example flow cytometry plots from HA-PLA-FITC BSA incubation with (A) U87 cells and (B) Jurkat cells at 4 □C. Flow cytometry data was analyzed over triplicates to calculate (A) the percent of cells positive for FITC-BSA and (B) the mean fluorescence intensity. * p<0.05, ** p<0.01, *** p<0.001, **** p<0.0001

**Figure 6.**
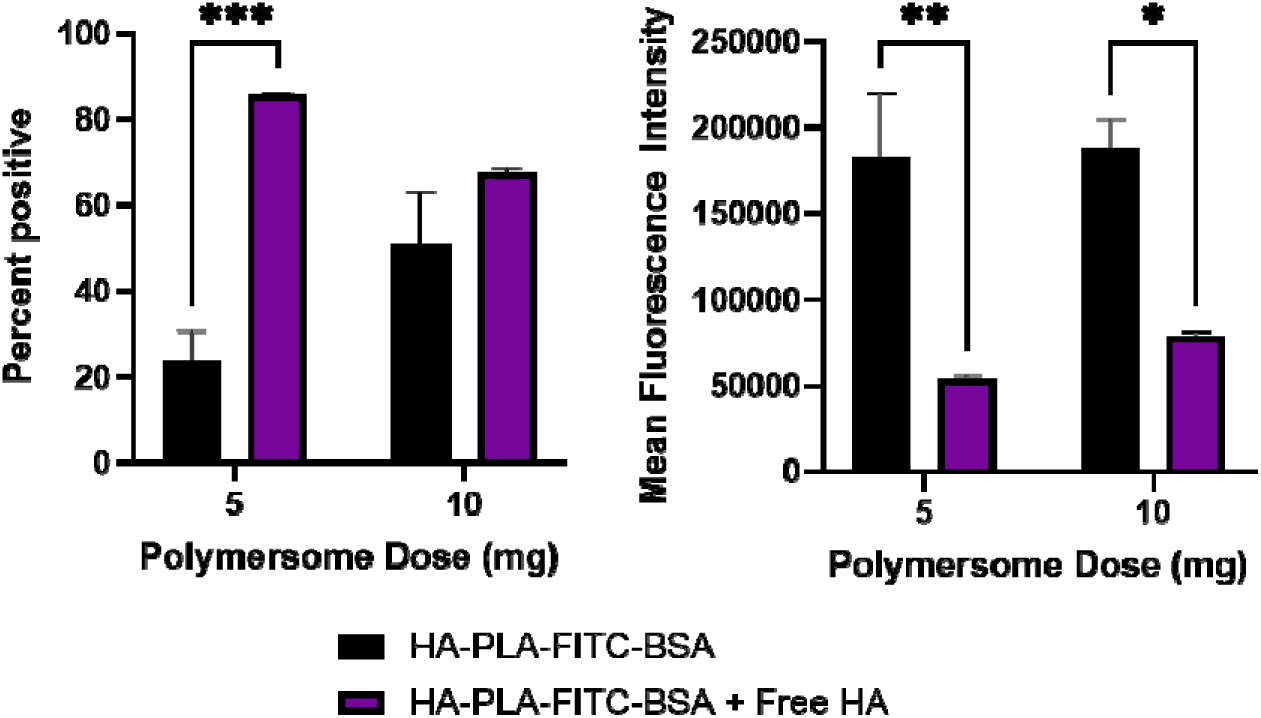
Flow cytometry data from competitive uptake experiments, where the addition of 10 mg/mL free HA (7 kDa) increased the number of positive cells (left) but dramatically decreased the overall mean fluorescence intensity (right).

This uptake mechanism has been confirmed previously for lipid nanoparticles (LNPs) conjugated with HA, with breast cancer being the most widely studied^39^. Specifically, Song et al. reported increased MFI for HA-functionalized LNPs loaded with DOX when administered to MCF-7 cells compared to HepG2 cells, due to increased CD44 expression in MCF-7 cells^40^. Mahira et al. quantified CD44 expression levels in PC-3 and DU-15 cells (prostate cancer) before and after dosing with HA-coated LNPs, where a decrease in CD44 expression was observed post-treatment, suggesting the receptors were used for internalization^41^. Shahriari, et. al confirmed that HA-polycaprolactone based polymersomes were internalized in CD44 expressing 4T1 and MCF-7 cell lines at higher amounts prior to the addition of free HA, also suggesting this same mechanism^42^.

### Free Doxorubicin performance

To determine the concentration of DOX required to induce U87 apoptosis, cells were incubated with free DOX at increasing concentrations for 5 days. After 24 hours, the cells were 75% viable, indicating the requirement of a longer duration to induce apoptosis. It required at least 3 days post incubation to observe a that reduced to 50% (Figure 3A). The viability data at 3,4, and 5 days post incubation of free DOX was fit to a curve to calculate the corresponding IC50 values, with values in the table reported as the 95% confidence interval (Figure 7). The best fit IC50 value of free DOX over these three days averaged to be 2.36 ± 0.23 µg/mL, indicating that our polymersome formulations with 6 and 24 µg DOX/mL, with loaded contents of 5.94 and 23.44 ± 0.45 µg DOX/mL respectively, should be sufficient to initiate apoptosis.

**Figure 7.**
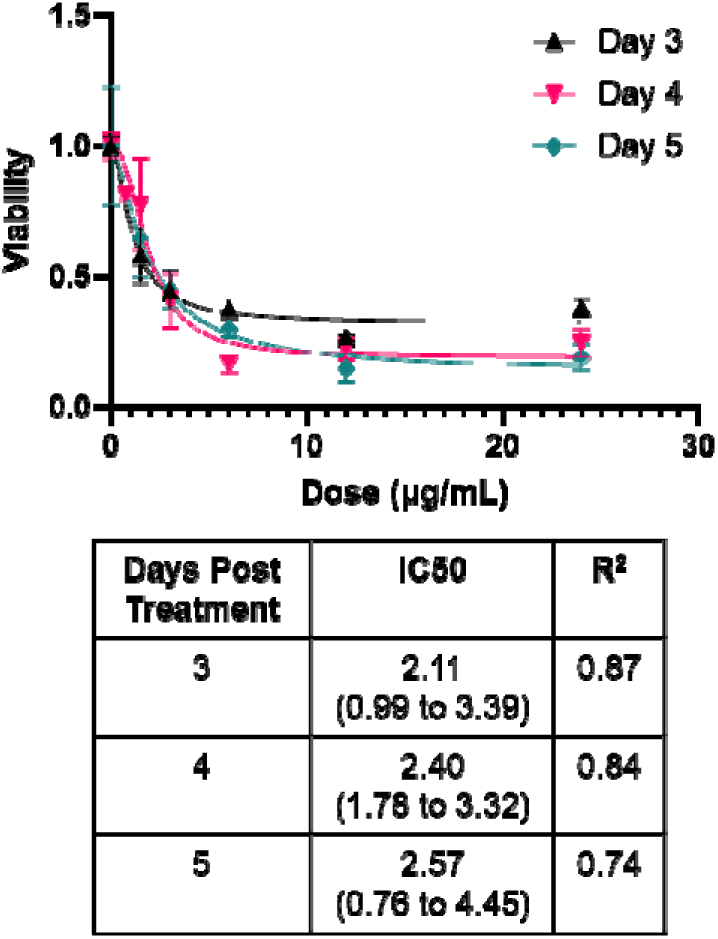
Viability of U87 cells over multiple doses of free DOX at 3, 4, and 5 days post-treatment is plotted to calculate the tabulated IC50 values.

### HA-PLA-DOX polymersomes reduce viability of U87 cells

The safety profile of the HA-PLA polymersomes on U87 cells was evaluated using MTS proliferation assay (Figure 8). U87 cells were treated with 6 and 24 µg/ml of HA-PLA-DOX polymersome system for 4 and 5 days, with empty polymersome and free DOX were included as controls. High cell viability of 90% or greater was observed when treated with empty HA-PLA polymersomes as expected, indicating that the polymersomes itself without DOX did not pose any cellular toxicity^12,43^. While free DOX was able to reach an IC50 at 5 days post incubation, neither HA-PLA-DOX formulation explored led to 50% viability or less. Notably, viability of U87 was still decreased when DOX was delivered via polymersome and the measured viability was not statistically different from free DOX at any time point. However, this increased viability could be due to the slower release of DOX from our polymersomes (Figure 2,3); it is known that at least 3 days are needed to reach IC50 of free DOX (Figure 7) and it is readily bioavailable in cells. An increased number of days may allow for IC50 values of HA-PLA polymersomes to be calculated. Similar behavior was observed when free DOX was compared to Doxil, an FDA-approved liposomal DOX formulation, with IC50 values of free DOX and Doxil calculated to be 0.25 µM and 0.7 µM, respectively^44^.

### HA-PLA-DOX induces inhibition of topoisomerase II in U87 cells

We treated U87 cells with HA-PLA-DOX system and extracted DNA followed by a test of topoisomerase activity using the TOPOII assay. After running gel electrophoresis (Figure 9) on the reaction products with catenated and linear DNA as positive controls, the treated DNA extract of both free DOX and HA-PLA-DOX formulations produced significantly brighter bands retained in the well, indicating that inhibition of TOPOII occurred. This confirms that HA-PLA-DOX uses the same therapeutic mechanism as free DOX. While multiple bands may be seen further down the gel, this was representative of multiple types of decatenation products (nicked and intact circular); both indicated that TOPO II-induced decatenation. Literature reports the effect of DOX on TOPO II inhibition through TOPO II assays, evaluating varying concentrations of DOX on kDNA cleavage patterns^45^. Complete inhibition would result in no products traveling down the length of the gel, indicating higher concentrations may be required for more significant results^45–47^.

**Figure 8.**
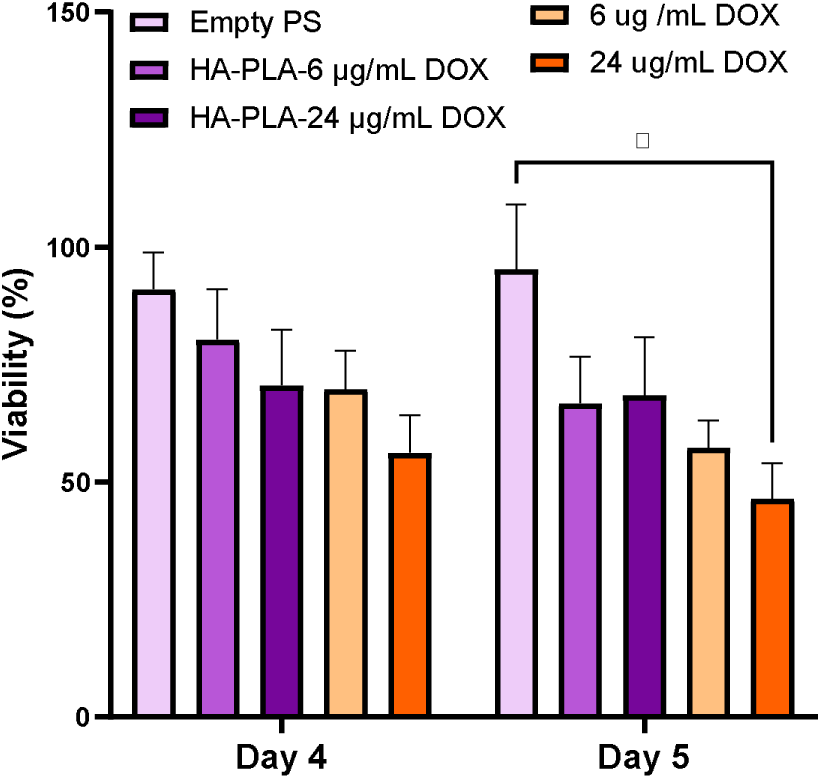
MTS Viability Assay was performed at 4 and 5 days post incubation with empty polymersomes (PS), various free DOX and HA-PLA DOX concentrations. *p<0.05

**Figure 9.**
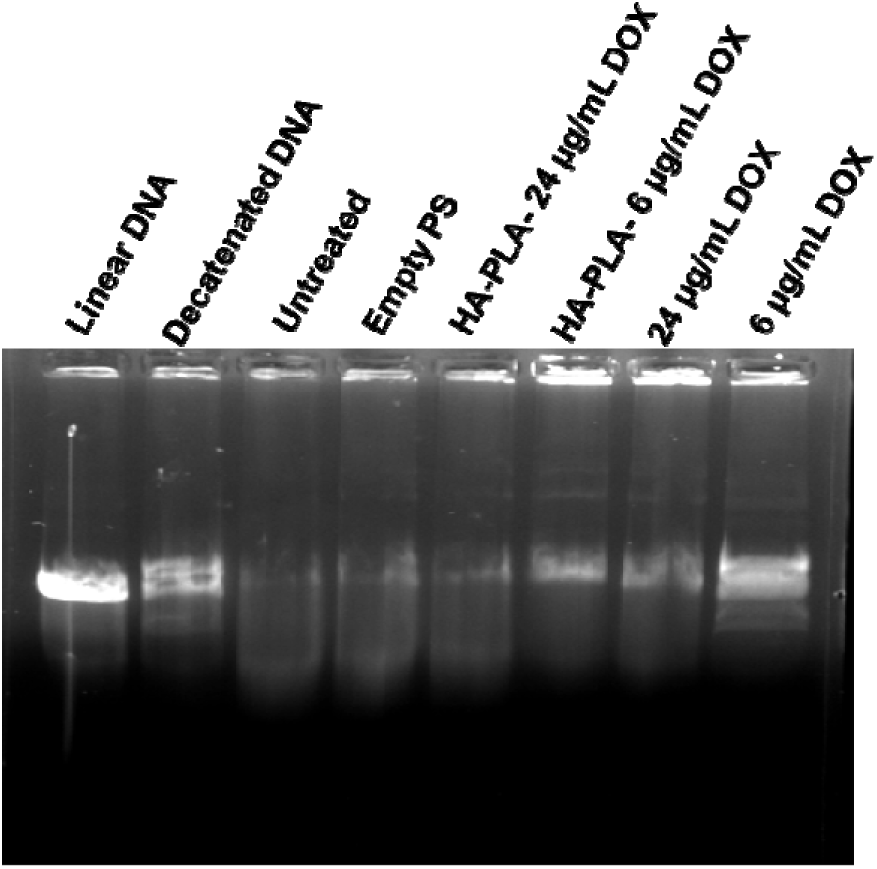
Ethidium bromide stained 1% agarose gel after electrophoresis to evaluate inhibition of TOPOII Activity. Lane 1: linear DNA marker (TOPOGen), Lane 2: decatenated DNA marker (TOPOGen). Lanes 3-8 contain DNA extracted from U87-MG cells 5 days after treatment. Lane 3: untreated, Lane 4: empty polymersome treated, Lane 5: 24 μg/mL DOX-loaded polymersomes, Lane 6: 6 μg/mL DOX-loaded polymersomes, Lane 7: 24 μg/mL free DOX, Lane 8: 6 μg/mL free DOX

## Conclusions

This study proposes a HA-PLA-DOX polymersome system for efficient delivery of chemotherapeutic agent doxorubicin in glioblastoma tumors. Polymersome-based delivery systems offer unique advantages for drug delivery, including specificity and biodegradability in human tissues. Several treatment strategies have not worked in glioblastoma due to constraints posed by the BBB. In summary, we have synthesized an HA-PLA copolymer that self-assembled into polymersomes through the solvent injection method. This method produced polymersomes with an average diameter of 101.8 ± 16.5 nm and ζ-potential of -11.3 ± 1.5 mV, characterized using TEM and DLS. These HA-PLA polymersomes achieved high encapsulation efficiencies of DOX above 98% when loading up to 24 µg DOX/mL. Our HA-PLA polymersomes were found to deliver DOX slowly over time, following a diffusion- and degradation-driven release model, with later time points favoring PLA hydrolysis in an environment mimicking the more acidic tumor microenvironment. Specifically, we find that HA-PLA polymersomes effectively infiltrate CD44-expressing glioblastoma cells via CD44 receptor-mediated endocytosis and effectively deliver DOX. We next confirmed that DOX-loaded polymersomes can reduce tumor cell proliferation in a human glioblastoma tumor cell line, U87, at 6 µg/mL and 24 µg/mL where they inhibit TOPOII. In this study, we observe efficient delivery of DOX through this nanoparticle system. Our work in an *in vitro* model of glioblastoma shows encouraging evidence of the potential for therapy. Subsequent work will focus on testing the efficacy of this system *in vivo* animal models of glioblastoma. Further, the ability of our HA-PLA polymersomes to cross the BBB will require ligand tagging to enable efficient trafficking and delivery at the tumor site. Overall, we expect this system to be efficient in glioblastoma through the CD44-mediated receptor endocytosis and acidic tumor microenvironment-mediated hydrolysis of the polymersome.

## Author contributions

**Molli Garifo:** Writing – original draft, Validation, Investigation, Formal analysis, Data curation, Conceptualization. **Apoorvi Chaudhri:** Methodology, Validation, Formal analysis, Investigation, Writing – original draft, Visualization. **Pranavi Thatravarthi:** Validation, Data curation. **Torrick Fletcher Jr.:** Validation, Data curation. **Jessica Larsen**: Writing – review & editing, Visualization, Supervision, Project administration, Funding acquisition, Conceptualization.

## Data availability

The data supporting this article have been included as part of the Supplementary Information

## Acknowledgements

We thank the Clemson Light Imaging facility for access to the confocal microscope. Molli Garifo was partially supported by US Department of Education GAANN #P200A210105. Apoorvi Chaudhri was supported in part by Postdoctoral Fellowship PF-24-1319605-01-IBCD from the American Cancer Society. TJ Fletcher was supported in part by NSF DMR # 1950557. Jessica Larsen was supported by NSF CAREER Award #2047697, NIH DP2NS148060, and P20GM103499.

