## Supplemental Tables and Figures for "Hyaluronic acid-b-polylactic acid polymersomes facilitate CD44-mediated delivery of doxorubicin to glioblastoma in vitro"

Apoorvi Chaudhri^1#^, Molli Garifo^1#^, Noah Arnold^1^, Jessica Larsen^1,2*^

^1^Department of Chemical and Biomolecular Engineering, Clemson University, Clemson, South Carolina, United States of America

^2^ Department of Bioengineering, Clemson University, Clemson, South Carolina, United States of America

^#^ These authors share equal contribution

Total number of tables: 1

Total number of figures: 1

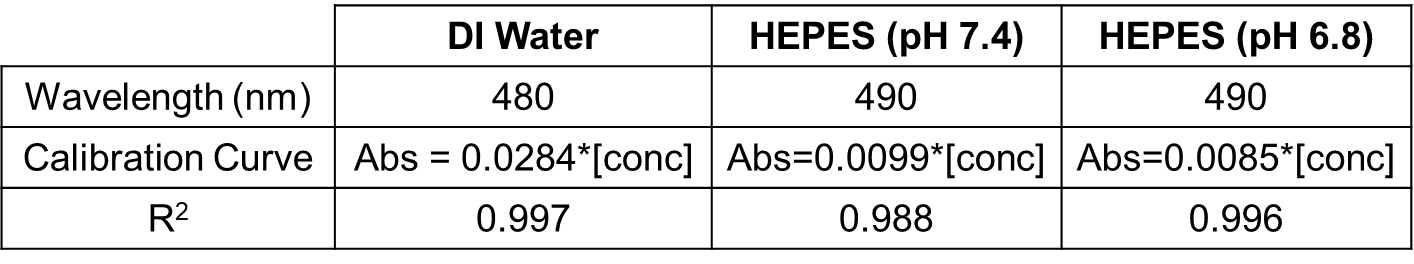

Table 1. Maximum absorbance wavelength from spectral scans ranging from 280 to 700 nm and example calibration curve data for known DOX concentrations in various buffers used in the release studies. Corresponding goodness-of-fit values are provided as R^2^.

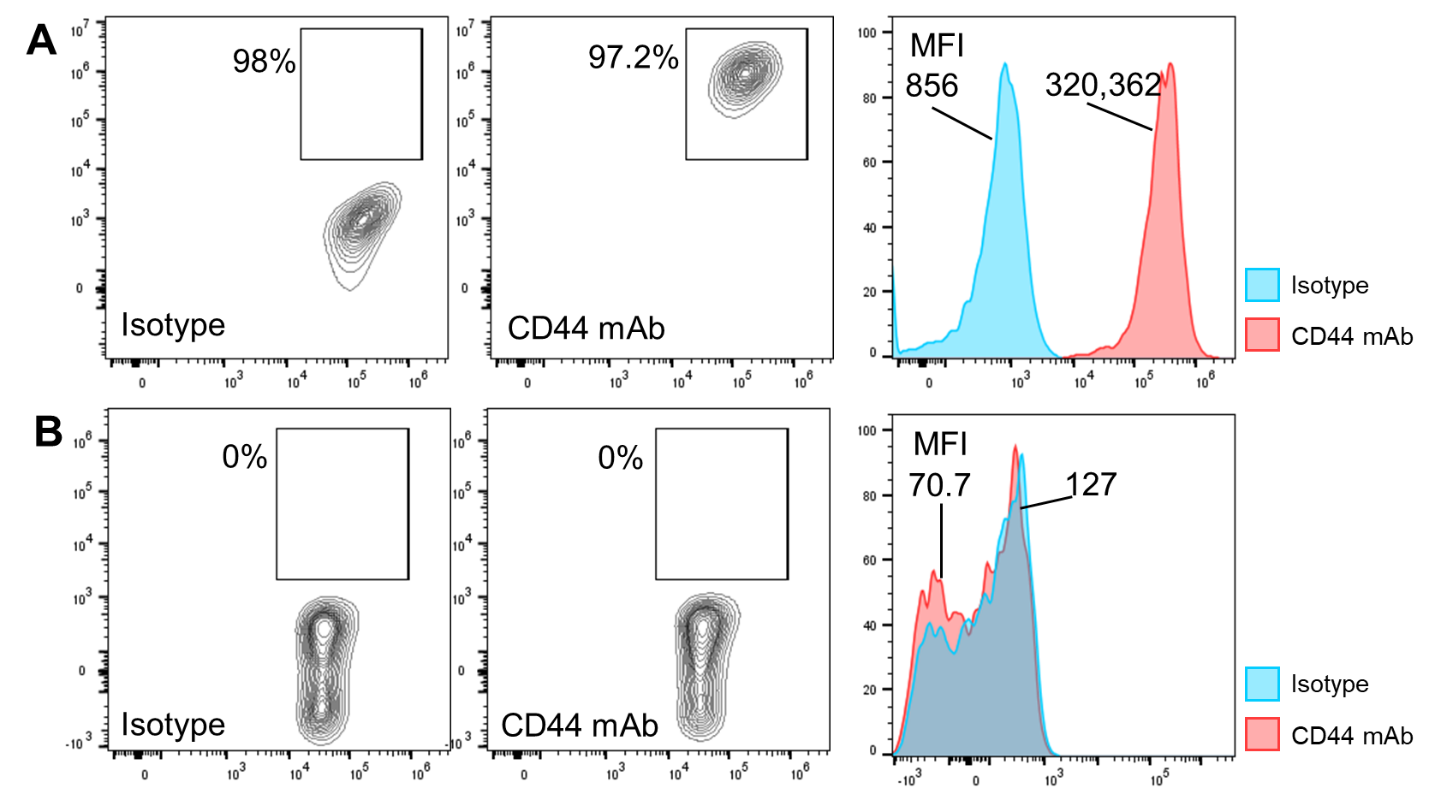

Figure 1. Flow cytometry data from (A)U87 and (B)Jurkat cells confirming the presence or absence of the CD44 receptor, respectfully, using CD44 mAb.
